# WNK kinase regulates plasma membrane levels of the WNT inhibitor RNF43

**DOI:** 10.1101/2025.10.08.681128

**Authors:** Gabriele Colozza, Ingrid Jordens, Eric Sosa, Jeongmin Ha, Szu-Hsien Sam Wu, Katherina Tavernini, Andrea Català-Bordes, Fiona Farnhammer, Noelia Urbán, Madelon M. Maurice, Bon-Kyoung Koo

## Abstract

The E3 ubiquitin ligases RNF43 and ZNRF3 are key negative regulators of canonical WNT signaling, promoting turnover of the WNT receptors FRIZZLED and LRP5/6 at the plasma membrane. While their mechanism of action is well established, how RNF43/ZNRF3 themselves are regulated remains unclear. Here, we identify WNK kinases as novel upstream regulators of RNF43 through proximity labeling proteomics. Using gain- and loss-of-function approaches, we show that WNKs control RNF43 surface localization and thereby its ability to ubiquitinate and downregulate WNT receptors. Pharmacological inhibition of WNKs increases RNF43 membrane abundance and enhances WNT suppression – an effect abolished in RNF43/ZNRF3 double knockout cells and organoids. Mechanistically, WNK inhibition alters RNF43 trafficking and ubiquitination, revealing a role for WNKs in regulating its plasma membrane distribution. These findings define a new regulatory axis linking the pro-WNT activity of WNKs to RNF43/ZNRF3-mediated feedback inhibition. Targeting WNK now offers a novel therapeutic strategy to restore WNT pathway control in cancers with RSPO fusions or RNF43 mutations.

## Introduction

The WNT/β-catenin signaling pathway (also known as canonical WNT) is conserved throughout metazoan evolution and plays a major role during embryonic development of multicellular animals (1, 2). In adult tissues, canonical WNT controls self-renewal and maintenance of tissue-specific stem cells, where optimal WNT signaling levels are required to keep the homeostatic balance between proliferation and differentiation. Dysregulation of WNT signals is at the basis of several diseases, including cancer (2, 3). For example, constitutive activation of WNT signaling is a key driver in the development and progression of colorectal cancer (CRC). WNT antagonists have evolved to prevent uncontrolled overstimulation; many of them are transcriptional targets of WNT itself, thereby forming a sophisticated genetic circuitry with multiple negative feedback loops. Among these inhibitors, RNF43 and its paralogue ZNRF3 (hereafter, R/Z) offer an excellent example of signaling regulation at the receptor level (1, 4). Both R/Z are transmembrane E3 ubiquitin ligases containing a Really Interesting New Gene (RING) domain, and target the WNT receptors FRIZZLED (FZD) and LRP5/6 for endolysosomal degradation in a ubiquitination-dependent manner (5, 6). By decreasing the cell surface levels of WNT receptors, R/Z efficiently dampen WNT signaling. Besides the RING domain, conferring the E3 ubiquitin ligase activity, R/Z also feature a Protease Associated (PA) domain, located on their extracellular portion (ectodomain) and involved in protein-protein interactions (7), a transmembrane domain (TM) which may dictate selectivity for FZD receptors (8), and a long cytoplasmic tail required for the regulation of R/Z activity (4).

Extracellularly, the secreted WNT agonists R-SPONDINS (RSPO)1/2/3/4 form a multiprotein complex with R/Z and the seven transmembrane receptors LGR4/5/6 (9–13). This complex is rapidly endocytosed, preventing FZD/LRPs receptor degradation by R/Z and thus strengthening WNT signaling. In other words, RSPO indirectly mediates an increase in the cell surface levels of WNT receptors that boosts signaling. A few studies also shed light on the intracellular machinery of R/Z, identifying key components that regulate its activity. For example, dishevelled (DVL), a core component involved in the transduction of WNT signals, interacts with RNF43 and bridges it to FZD, promoting receptor ubiquitination and degradation (14). This interaction requires a specific DVL binding region present on the cytoplasmic tail of RNF43. In addition, RNF43 contains a cytoplasmic phospho-switch for its suppressive role on FZD (15). More recently, we have uncovered an unexpected link between DAAM1/2 and RNF43. In fact, DAAM1/2 are required for FZD internalization following ubiquitination by RNF43, ensuring efficient clearance of WNT receptors from the cell surface (16).

In an effort to better characterize the molecular machinery underlying R/Z mechanistic regulation, we performed a high-throughput interactome analysis based on APEX2-mediated protein proximity labeling (17, 18), to identify novel modulators of RNF43. As a result of our work, we present here evidence that the With-NO-Lysine [K] (WNK) kinases are previously unrecognized regulators of RNF43. WNKs are a unique family of serine/threonine protein kinases distinguished by the atypical placement of a catalytic lysine within their kinase domain (19). They play pivotal roles in regulating ion transport processes, particularly in the kidney. Mutations in WNK1 or WNK4 have been implicated in conditions such as Gordon’s syndrome, characterized by hypertension and hyperkalemia (19). Beyond their established functions in ion homeostasis, WNK kinases have garnered attention for their involvement in oncogenesis. Dysregulation of WNKs has been linked to tumor growth, metastasis and angiogenesis through complex mechanisms (20). For instance, WNK1 has been identified as a gene uniquely associated with a subset of invasive cancers, suggesting its potential as a therapeutic target (21).

Intriguingly, WNK kinases intersect with the WNT signaling pathway. In the fruit fly *D. melanogaster*, studies have shown that WNK kinases can modulate canonical WNT by influencing the phosphorylation state of Dishevelled (22, 23). Furthermore, in mammalian cells, WNK is required for β-catenin stabilization (24). However, despite these initial findings, their role in WNT signaling regulation remains elusive without clearly known mechanisms. In summary, our present work shows that WNK regulates membrane trafficking of RNF43, and inhibition of WNK increases RNF43 levels on the plasma membrane and, consequently, leads to WNT inhibition.

## Results

### A dual proximity labeling strategy to probe RNF43 interactomes

We reasoned that a large-scale approach aimed at identifying RNF43 protein-protein interactions would provide insight into its regulatory mechanisms. To this aim, we relied on two promiscuous biotin labeling enzymes, the genetically engineered soybean ascorbate peroxidase, APEX2 (17, 18), and horse radish peroxidase (HRP) (25). Mass-spectrometry (MS)-coupled, proximity-dependent biotin labeling has emerged as a widely used tool to study protein-interaction networks (26). Both APEX2 and HRP have been successfully applied in numerous studies to characterize subcellular proteomes and signaling protein complexes, including those involved in WNT pathway regulation (27–29).

Using RNF43 as a bait, we generated two different fusion constructs: in the first, we appended the coding sequence for APEX2 at the C-terminal of its cytosolic tail **(Fig. 1a)**; in the second, we inserted the cDNA encoding for HRP within the RNF43 ectodomain, immediately after the signal peptide required for RNF43 plasma membrane localization **(Fig. 1a)**. This dual strategy would allow biotinylation of RNF43 interactors on both the cytoplasmic and extracellular sides of the plasma membrane, where APEX2 and HRP are most effective, respectively. When overexpressed in HEK293T cells, both constructs retained their ability to degrade FZD receptor and inhibit the WNT-responsive TOPFlash luciferase reporter **(Fig. 1b, c)**, with effects comparable to wild type (WT) RNF43. This indicated that neither the APEX2 nor HRP tags interfered with RNF43 biological function. The activity of these fusion proteins was further validated *in vivo* using a classical embryological assay in *Xenopus laevis* embryos **(Fig. 1d)**. Injection of *xWnt8* mRNA into the ventral blastomere of 8-cell stage embryos induced secondary axis formation, as expected (30) **(Fig. 1d, e)**. Co-injection of either *RNF43-APEX2* or *HRP-RNF43* mRNAs inhibited this phenotype, similar to WT RNF43, confirming the constructs functional integrity **(Fig. 1d, e)**.

**Figure 1.**
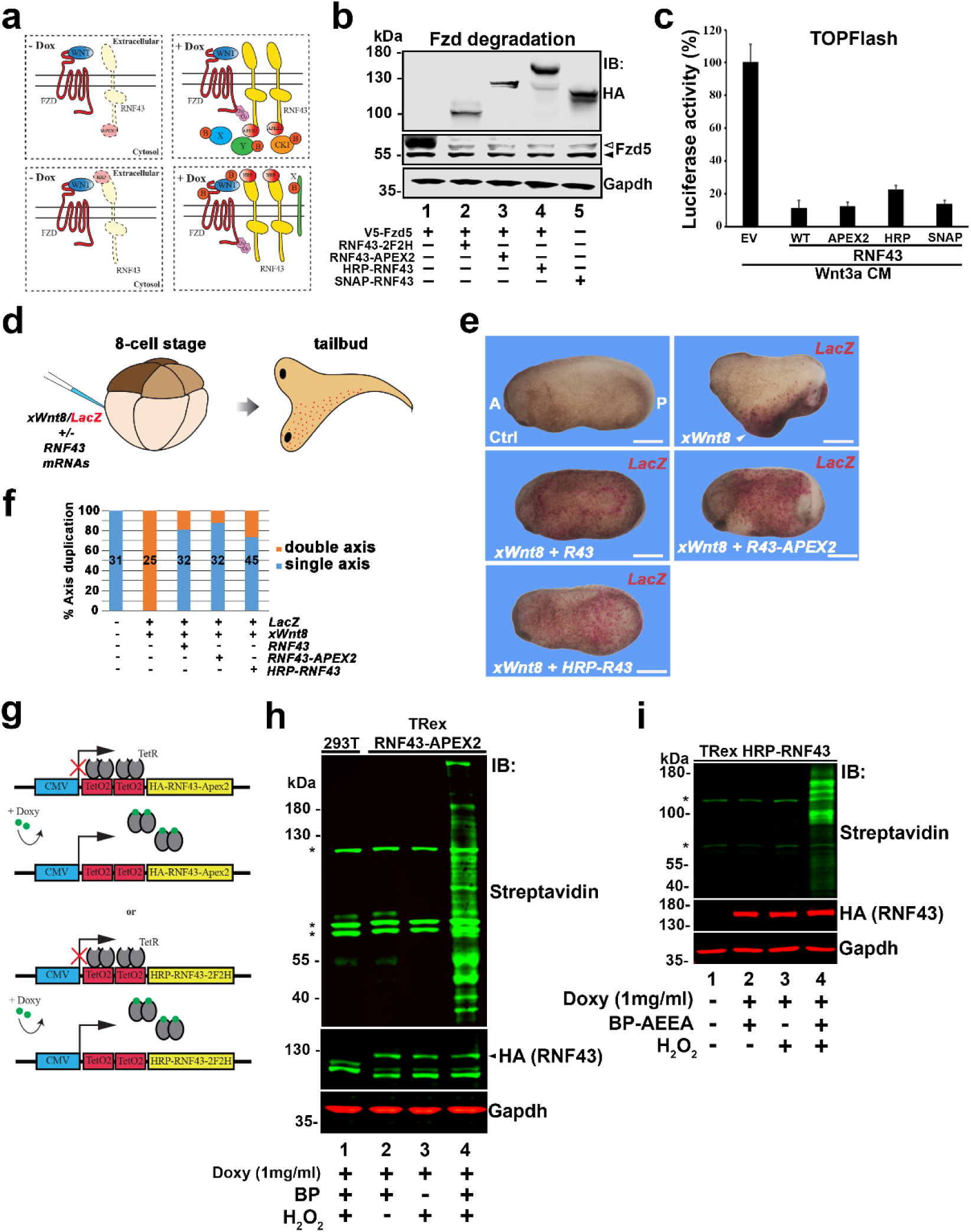
Generation of stable cell lines expressing HRP-RNF43 and RNF43-APEX2 chimeric constructs. **a** Schematic showing HRP- and APEX2-tagged RNF43 constructs and the different cell compartments where the biotinylating activity occurs. **b** Western blot showing degradation of overexpressed mouse Fzd5 protein by the indicated RNF43 constructs, in HEK 293T cells. Note that RNF43 chimeras retain comparable activity to WT RNF43. **c** Topflash luciferase assays in HEK293T cells showing comparable Wnt signaling inhibition by overexpressed WT and chimeric RNF43 DNA constructs. **d, e** double axis assay in *Xenopus* embryos. Injections of Wnt8 mRNA in a single ventral blastomere induces the formation of a secondary axis (white arrowhead, top right panel in **e**), which is promptly inhibited by co-injection of the indicated *RNF43* mRNAs. Scale bars equal to 500 μm. **f** Quantification of the experiment shown in **e**. **g** schematic showing the doxycycline inducible constructs containing HRP-RNF43 and RNF43-APEX2, used to generate stable cell lines. **h, i** Western blots showing biotinylating activity by doxy-induced RNF43. Note that biotinylation only occurs in samples treated with biotin-phenol (BP) and H_2_O_2_.

Next, we established two stable transgenic cell lines by transfecting T-REx 293 cells with plasmids encoding RNF43-APEX2 or HRP-RNF43, under the control of two tetracycline operator 2 (TetO_2_) sites downstream of a strong cytomegalovirus (CMV) promoter. In this inducible system, expression of the RNF43 constructs is repressed by the presence of the constitutive Tet Repressor (TetR) protein and activated upon addition of doxycycline (dox) to the culture medium **(Fig. 1g)**. Western blot (WB) analysis of cell extracts from monoclonal transgenic lines confirmed robust induction of RNF43 fusion proteins following overnight dox treatment **(Fig. 1h, i**, lanes 2–4). Furthermore, strong biotinylation activity was detected upon co-incubation with biotin-phenol (BP) and H_2_O_2_ in dox-induced cells, as shown by WB **(Fig. 1h, i**, lane 4) and immunofluorescence (IF) analyses **(Fig. S1a, b)**. For simplicity, we refer to these cell lines as T-REx RNF43-APEX2 (TRA) and T-REx HRP-RNF43 (THR).

Both TRA and THR cells maintained stable, inducible expression of RNF43 fusions and biotinylation activity even after multiple passages **(Fig. S2a, b)**. Upon induction with dox, both constructs strongly inhibited WNT signaling, as confirmed by WB assays showing consistent suppression of unphosphorylated β-catenin and phospho-Dvl2 **(Fig. S2c-e)**, two well-established markers of active WNT signaling. Importantly, both RNF43-APEX2 and HRP-RNF43 remained responsive to the secreted WNT agonists R-spondin 1 and 2 (Rspo1/2), which derepressed WNT signaling even under conditions of RNF43 overexpression **(Fig. S2c, d)**.

### Proteomics analysis of RNF43 interactome revealed WNK1 and SORTILIN as novel interactors

Using the cell lines described above, we set out to identify RNF43-interacting proteins by performing large-scale proximity labeling followed by mass spectrometry analysis **(Fig. 2a** and **Fig. S3a, b)**. For the labeling reaction, TRA cells were incubated with biotin-phenol (BP), which diffuses across the plasma membrane into the cytosol. In contrast, THR cells were treated with the membrane-impermeant analog biotin-AEEA-phenol (BP-AEEA), thereby restricting labeling to the cell surface. Proteomic analysis of the purified biotinylated proteins revealed several noteworthy findings. Volcano plots and heatmaps show that HRP-RNF43-mediated biotinylation predominantly captured transmembrane receptors and secreted ligands, including WNT5a and Rspo1 **(Fig. 2b, c** and **Supplementary Table 1)**. Among the transmembrane proteins, we identified several known RNF43 targets, such as the non-canonical WNT co-receptors PTK7 and ROR1/2, the canonical WNT co-receptor LRP6, and multiple Frizzled (FZD) family members (5, 6, 31). We also detected components of the TGF-β/BMP and EGF/FGF receptor families, which have recently been implicated as substrates of the RNF43/ZNRF3 axis (32, 33) **(Fig. 2b, c** and **Fig. S4a)**.

**Figure 2.**
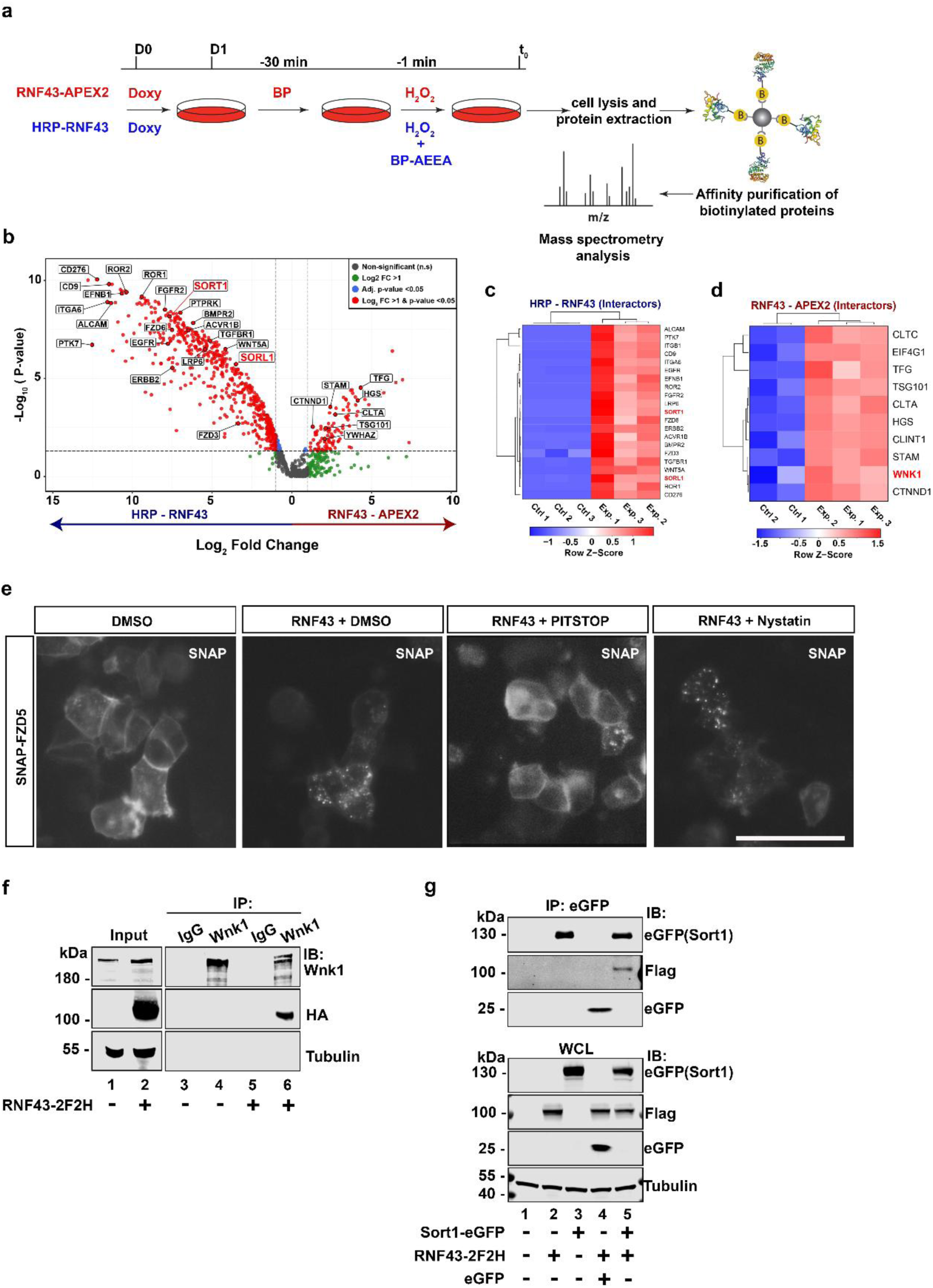
Identification of RNF43 binding partners via proximity labeling-coupled mass spectrometry. **a** Schematic of the biotinylation experiment coupled to Mass spectrometry analysis. **b** Volcano plot illustrating logarthmic fold changes of proteins identified in HRP- and APEX2-mediated biotinylation experiments. A total of 577 proteins (HRP-RNF43) and 183 proteins (RNF43-APEX2) met the threshold for statistical significance (≥ 2-fold change, *p* ≤ 0.05) and are denoted in red. Proteins with a ≥ 2-fold change but not statistically significant are shown in green, while proteins with statistically significant *p*-values (*p* ≤ 0.05) but fold change < 2 are shown in blue. Fold changes and *p*-values were derived from triplicate experiments and corresponding controls using imputed data. **c** Heatmap of proteomic expression of HRP-RNF43 biotinylation interactors. Row z-scores represent the number of standard deviations each protein’s expression deviates from its mean across triplicate experiments. Red and blue indicate higher and lower expression levels, respectively. Differential expression of 21 HRP-RNF43 interactors is shown, with SORT1 and SORL1 highlighted in red text. **d** Heatmap of proteomic expression of 10 RNF43-APEX2 interactors based on duplicate control and triplicate experimental conditions. The interactor WNK1 is highlighted in red text. **e** SNAP-labeling of SNAP-Fzd5 expressing cells. In cells co-transfected with RNF43, Fzd5 rapidly relocates into cytosolic vesicles (puncta). However, treatment with the clathrin inhibitor PITSTOP partially suppresses Fzd5 endocytosis. Nystatin, an inhibitor of caveolin-dependent endocytosis, has no effect on RNF43-dependent Fzd5 endocytosis. Scale bar represents 50 μm. **f, g** Immunoprecipitation and Western blot analysis showing interactions between RNF43 and endogenous WNK1 and overexpressed SORTILIN. Unrelated IgG (**f**) and eGFP (**g**) were used as negative controls.

Conversely, APEX2-mediated labeling of RNF43 predominantly enriched proteins associated with endocytosis and endosomal trafficking **(Fig. 2b, d** and **Fig. S4b)**, as well as components of the ER/Golgi secretory pathway and various cytosolic kinases **(Supplementary Table 2)**. The identification of endosome-resident proteins was expected, as RNF43 is known to direct FZD and other substrates toward endo-lysosomal degradation (5, 6). Notably, proteins involved in clathrin-mediated endocytosis (CME) were significantly enriched, suggesting that this pathway plays a prominent role in RNF43-driven cargo internalization and degradation. Supporting this idea, inhibition of CME with Pitstop-2 strongly impaired RNF43-mediated internalization of fluorescent SNAP-labeled FZD, whereas the caveolin inhibitor Nystatin had no effect **(Fig. 2e)**.

Interestingly, the two proteomic datasets showed minimal overlap. This may be explained by the fact that overexpressed RNF43 predominantly localizes to intracellular vesicles—including the ER, Golgi, and endosomes—leading the APEX2-based intracellular labeling to favor proteins in these compartments. In contrast, the use of membrane-impermeant BP-AEEA and an extracellular-facing HRP prevented saturation by the dense proteome of internal membranes, promoting the labeling of proteins on the cell surface. It is also possible that the majority of RNF43 targets and regulatory interactions occur at the plasma membrane, which represents only a minor fraction in the cytosolic labeling context. In agreement with the latter hypothesis, the HRP-RNF43 proteome provided greater insight into the regulatory networks linking RNF43 to cancer-associated signaling pathways **(Fig. S4c, d)**. UpSet analysis revealed that a substantial subset of HRP-RNF43-enriched proteins were shared across multiple malignancies, including gastric adenocarcinoma, neuroblastic tumors, and tumor-associated vasculature **(Fig. S4c)**, suggesting a convergence of RNF43-associated signaling mechanisms in oncogenic contexts. Gene-concept network analysis further demonstrated that several HRP-RNF43 interactors were central to multiple cancer-related pathways, with proteins such as ERBB2, FGFR2, and CD276 serving as hubs connecting diverse malignancies **(Fig. S4d)**. These findings highlight the broad relevance of RNF43-interacting proteins in tumorigenic signaling networks and support a model in which RNF43 modulates key effectors implicated in multiple cancer subtypes. Despite these differences, both labeling approaches successfully identified novel RNF43-associated proteins. Notably, the cytosolic kinase WNK1 (from APEX2 labeling) and the transmembrane sorting receptor SORTILIN (34) (from HRP labeling) emerged as new candidates **(Fig. 2b–d)**. Co-immunoprecipitation assays confirmed robust interactions between RNF43 and endogenous WNK1 and transfected WNK2 **(Fig. 2f** and **Fig. S4e, f)**, as well as with SORTILIN and its homolog SORL1 **(Fig. 2g** and **Fig. S4g)**. Importantly, neither WNK nor SORTILIN protein levels were affected by RNF43 over-expression, suggesting they are unlikely to be direct targets of RNF43. Given previous evidence implicating WNK kinases in WNT signaling (22–24), though their precise role remains unclear, we decided to focus on their functional characterization.

### WNK regulates WNT signaling through RNF43

WNKs are serine/threonine kinases that play a major role in osmosensing and cell volume by regulating the trafficking and activity of several transmembrane transporters, like cathion-cloride transporters (CCC) (19). It is generally assumed that WNKs do not phosphorylate directly their effector proteins, but operate through a cascade involving the phosphorylation/activation of the intermediary kinases OSR1/SPAK (19) (**Fig. S5a**), which in turn phosphorylate the final targets of the WNK cascade. To determine the functional role of WNK in WNT signaling, we carried out a set of gain and loss of function experiments. In agreement with previous observations (22, 24), transfection of WNK2 was sufficient to increase WNT/β-catenin signaling in HEK293T cells, as determined by TOPFlash luciferase assay **(Fig. 3a)**. The increase in reporter activity was dependent on WNT ligand secretion, as the porcupine inhibitor IWP2 abolished the effects of WNK overexpression **(Fig. 3a)**.

**Figure 3.**
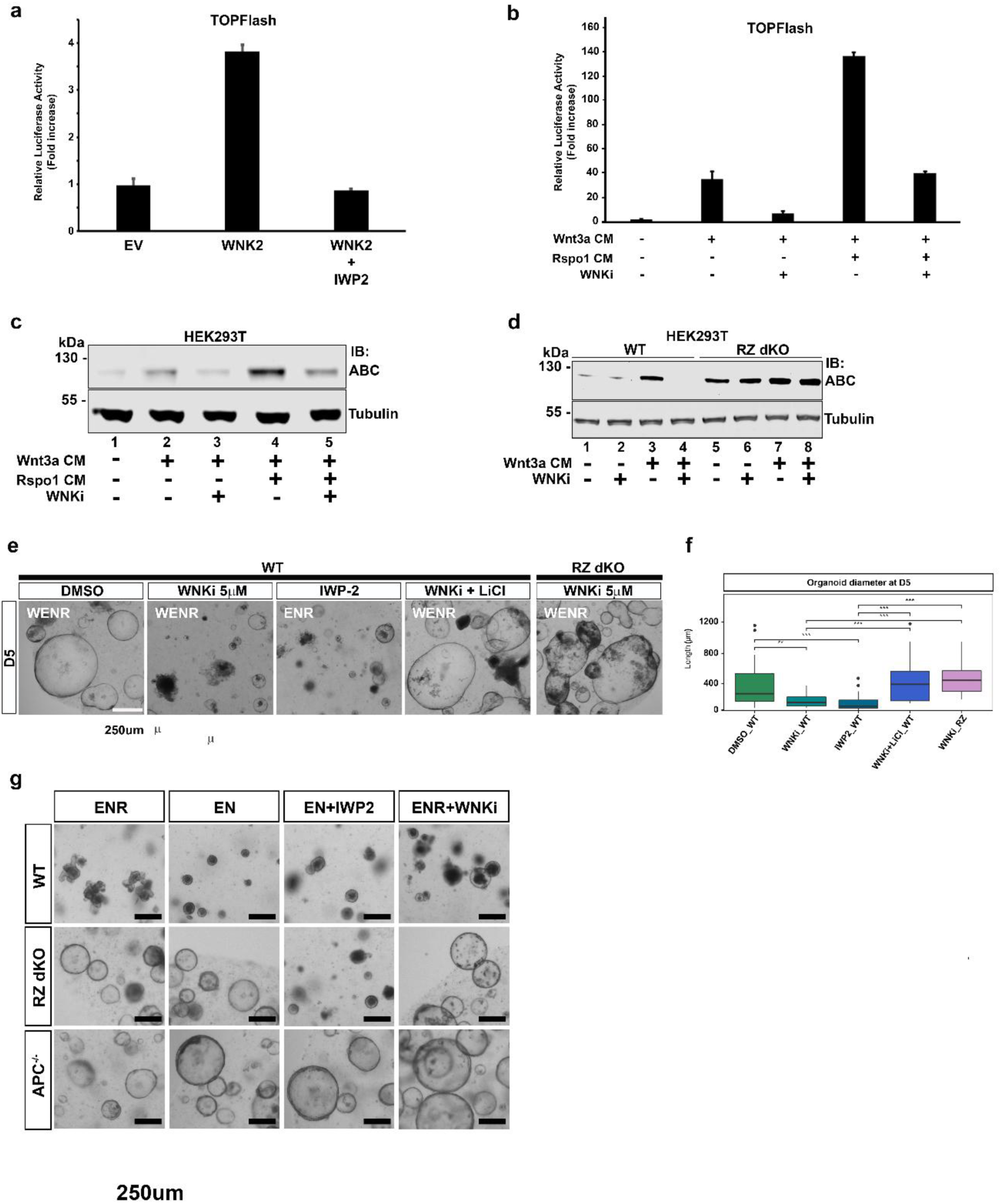
WNK inhibitor downregulates WNT signaling in a RNF43-dependent manner. **a** Overexpression of WNK2 increases WNT signaling, as measured via a TOPFlash reporter assay in HEK293T cells. WNT stimulation by WNK2 occurs at the ligand/receptor level, as inhibition of endogenous WNT secretion by IWP2 prevents WNK-mediated stimulation. **b** TOPFlash reporter analysis showing WNT signaling inhibition by the WNK inhibitor STOCK2S-26016 (WNKi for brevity). WNKi completely abolishes reporter activation by WNT3a ligand and reduces WNT boosting by R-spondin agonists. **c** Western blot of protein lysates from HEK293T cells stimulated with Wnt3a and Rspo1 conditioned media. Active (non-phosphorylated) β-catenin protein levels (ABC) are strongly reduced in the presence of WNKi. Non-phosphorylated β-catenin is a hallmark of active WNT signaling. **d** Western blot comparing ABC protein reduction by WNKi in WT HEK293T and RNF43/ZNRF3 double knock-out (RZ dKO) cells. Note that WNKi has no effect on ABC levels in the absence of RZ. **e** WNKi inhibits small intestinal organoid growth, similar to the well-known WNT inhibitor IWP2. Activation of WNT signaling by LiCl rescues growth in WNKi-treated intestinal organoids. Note that RZ dKO organoids are not affected by WNKi. Scale bar represents 250 μm. **f** Quantification of the experiment shown in e (n=3). **g** Small intestinal organoids from different genetic background respond differently to WNKi. Neither RZ dKO nor APC^−/-^ organoids are affected by WNKi. Scale bar represents 250 μm.

Given the availability of well-established soluble inhibitors that specifically interfere with WNK–OSR1/SPAK activation, we opted for a pharmacological loss-of-function approach rather than a genetic knock-out strategy. This decision was primarily driven by the presence of four WNK homologues (WNK1–4) in mammals, which complicates genetic targeting due to potential functional redundancy and compensatory mechanisms, while small molecule inhibitors have shown to possess broad but specific activity against all four WNK kinases. As expected, treating cultured cells with one of these inhibitors, namely STOCK2S-26016 (hereafter, WNKi) (35), potently inhibited signaling activation by the canonical WNT ligand, WNT3a, as shown in both WB and TOPFlash reporter assays **(Fig. 3b, c)**. Addition of Rspo1 conditioned medium (CM) greatly boosted WNT/β-catenin signaling, but its effects were strongly diminished in presence of WNKi **(Fig. 3b, c)**.

To better pinpoint the level at which WNK regulates the WNT pathway, we conducted epistatic analyses using TOPFlash reporter assays in HEK293T cells. Transfection with β-catenin expression constructs or treatment with GSK3 inhibitors robustly activated the WNT reporter, and these effects were largely unaffected by WNK inhibition **(Fig. S5b)**. These findings suggest that WNK acts upstream of β-catenin stabilization and GSK3 inhibition within the WNT signaling cascade. Interestingly, genetic double knock-out of RN43/ZNRF3 (RZ dKO) (36) or overexpression of a dominant negative form of RNF43 (5) rendered cells insensitive to WNK inhibition, while maintaining a strong response to WNT pathway activation **(Fig. 3d** and **Fig. S5b)**. Together with our previous observation that WNT activation by overexpressed WNK2 requires WNT ligand secretion **(Fig. 3a)**, these results suggest that WNK regulates WNT signaling at the level of ligand/receptor activation, likely through RNF43. Experiments conducted on mouse small intestinal organoids further corroborated these findings. Treatment with the WNK inhibitor (WNKi) significantly impaired the growth of cystic organoids cultured in WNT-enriched medium (WENR), compared to untreated controls, with effects comparable to those observed upon WNT inhibition using the porcupine inhibitor IWP2 **(Fig. 3e, f)**. Notably, organoid growth could be rescued by LiCl, a strong GSK3 inhibitor known to activate WNT signaling (1) **(Fig. 3e, f)**. In contrast to WT, RZ dKO organoids were resistant to WNK inhibition, as were APC^−/-^ organoids **(Fig. 3e–g)**. Altogether, these results indicate that RNF43/ZNRF3 act downstream of WNK and are epistatic to WNK in the regulation of WNT signaling.

### WNK regulates RNF43 cell surface levels

Regulation of signaling receptor abundance at the plasma membrane represents a critical control node for many signaling pathways. RNF43 modulates WNT signaling by controlling the cell surface levels of the WNT receptor Frizzled and LRP5/6, as well as non-canonical WNT receptors (5, 6, 31). However, emerging evidence indicates that RNF43 activity itself is tightly regulated through modulation of its own localization and abundance at the plasma membrane, a process that requires the interplay of different PTMs including phosphorylation and ubiquitination (15, 37, 38). Furthermore, WNK kinases are known to regulate trafficking and plasma membrane localization of several ion channels (19). Thus, we hypothesized that WNK could control WNT signaling by modulating the cell surface levels of RNF43. To test this possibility, we first overexpressed RNF43 constructs harboring a SNAP-tag on their ectodomain in HEK 293T cells. This modification, which did not interfere with RNF43 functions **(Fig. 1b, c** and **Fig. S2f),** allowed us to label specifically cell-surface RNF43 with a fluorescent molecule and follow its subcellular trafficking **(Fig. 4a)**. Cells treated with WNKi showed substantially higher levels of RNF43 on the plasma membrane, as compared to DMSO controls. Furthermore, while control cells showed many RNF43^+^ intracellular puncta 30’ after fluorescent labeling, these puncta appeared to be less prominent and closer to the membrane of WNKi treated cells **(Fig. 4a)**.

**Figure 4.**
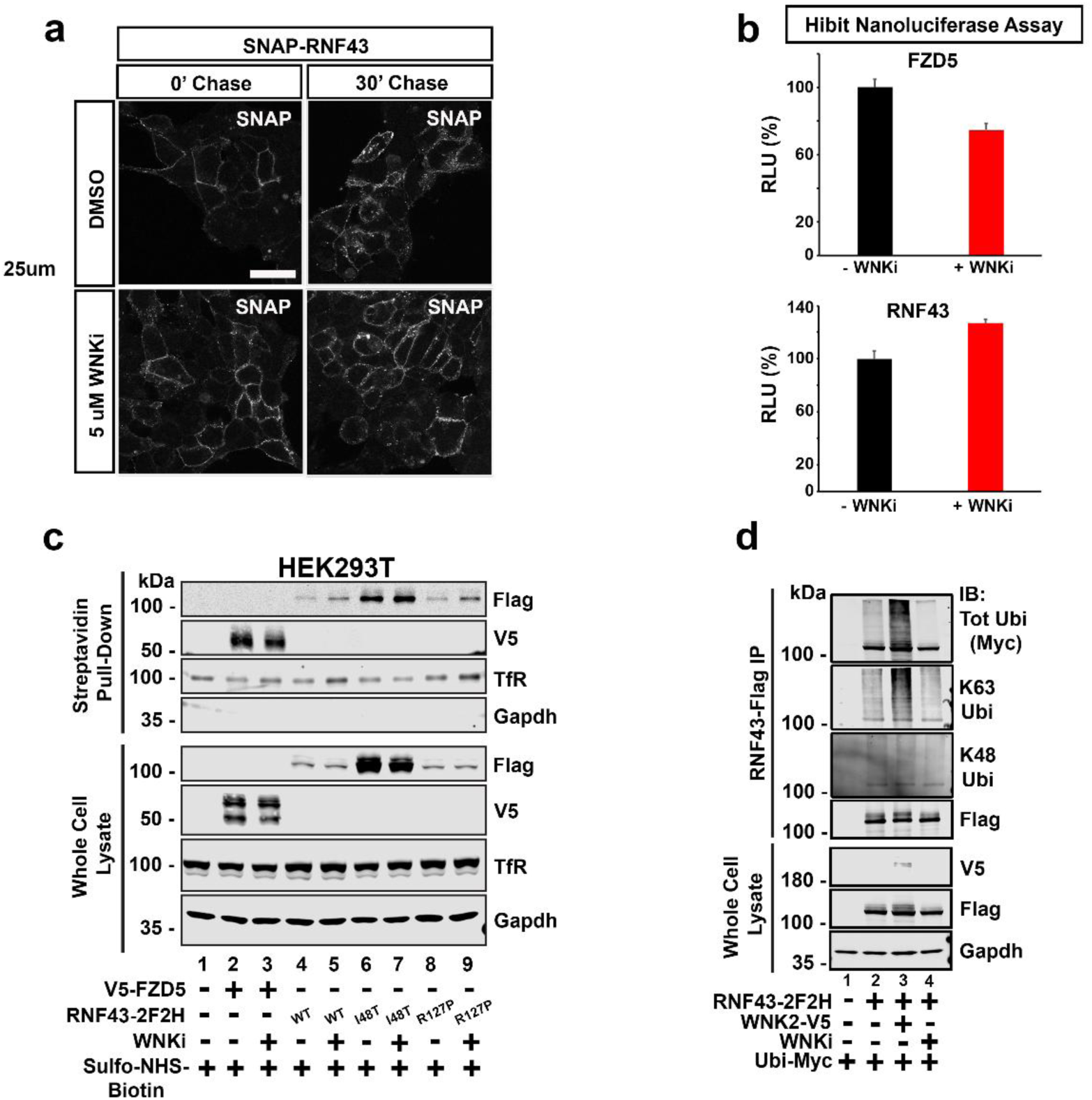
WNK inhibitor downregulates WNT signaling in a RNF43-dependent manner. **a** SNAP labeling in HEK293T cells transiently transfected with SNAP-RNF43. Treatment with WNKi increases cell surface levels of RNF43, most notably after 30’ chase. Scale bar equals to 20 μm. **b** HiBit nanoluciferase assays on HEK293T cells expressing HibiT-FZD5 or HiBit-RNF43. Luciferase activity measurements denote a decrease in the surface levels of FZD5 upon treatment with WNKi. Conversely, WNKi treatment increases RNF43 cell membrane levels. (n=3). **c** Cell surface protein biotinylation in HEK293T cells expressing the indicated RNF43 constructs. Western blot analysis confirms that WNKi increases the plasma membrane levels of RNF43. **d** RNF43 ubiquitination assay. When co-expressed with WNK2, RNF43 K63 ubiquitination levels increase, while K48 do not.

To precisely quantify the cell surface levels of RNF43, we employed the HiBiT/nanoluciferase reporter system. In this setup, overexpressed RNF43 is tagged with the HiBiT peptide on its ectodomain, which is exposed to the extracellular environment. Upon addition of the extracellular nanoluciferase component (LgBiT), the two fragments associate to reconstitute a functional luciferase, producing a luminescent signal. Since the complementation occurs exclusively at the cell surface, the intensity of the luminescence is directly proportional to the amount of RNF43 localized at the plasma membrane. As shown in **Fig. 4b**, WNK inhibition led to a 30% increase in RNF43 surface levels. HEK293 cells are known to express low levels of endogenous RNF43/ZNRF3 (6). If WNK negatively regulates RNF43 plasma membrane levels, then we would expect WNK inhibition to enhance RNF43 activity, leading to a reduction in the surface levels of its direct targets. Consistently, surface levels of HiBiT-tagged FZD5 were decreased upon WNK inhibitor treatment, supporting the notion of increased RNF43 activity at the plasma membrane **(Fig. 4b)**. To further validate the link between WNK activity and RNF43 localization, we performed cell surface protein biotinylation followed by pull-down and Western blot analysis. Consistent with our previous observations, WNK inhibition increased the amount of wild-type RNF43 at the plasma membrane, and reduced FZD5 **(Fig. 4c)**. Notably, we also detected increased surface levels of RNF43 I48T and R127P mutants, which are thought to accumulate in the endoplasmic reticulum (7) **(Fig. 4c)**. Taken together, our results suggest that WNK negatively controls the trafficking or stabilization of RNF43 at the cell surface.

Since ubiquitination has been shown to play a key role in regulating RNF43 plasma membrane localization and stability via endocytosis (37, 38), we tested whether WNK influences RNF43 ubiquitination. As shown in **Fig. 4d**, immunoprecipitation followed by Western blot analysis revealed that WNK overexpression increased RNF43 ubiquitination levels, particularly K63-linked ubiquitination, which is typically associated with the internalization of membrane proteins (39). Conversely, treatment with WNK inhibitor reduced K63-linked ubiquitination of RNF43 (**Fig. 4d**). Neither WNK overexpression nor inhibition significantly affected K48-linked ubiquitination. We did not observe major changes in total RNF43 protein levels, suggesting that ubiquitinated RNF43 may follow a fate other than lysosomal degradation – possibly recycling through endosomal compartments. Altogether, these results suggest that WNK promotes RNF43 internalization through K63-linked ubiquitination, regulating its membrane availability and consequently its anti-WNT activity.

### WNK inhibition compromises the growth of WNT-addicted organoids driven by RSPO fusions

To evaluate the potential of WNK inhibition as a therapeutic strategy for WNT-addicted tumors driven by chromosome fusions and rearrangements involving R-spondins (40), we established a mouse intestinal organoid model carrying a Ptprk–Rspo3 (PR) fusion (41). As shown by immunoblotting, PR fusion organoids express endogenous Rspo3 protein, in contrast to wild-type (WT) organoids, which lack Rspo3 expression and thus rely on exogenous Rspo supplementation for growth (**Fig. 5a**). Accordingly, PR fusion organoids were able to grow in Wnt3a/EGF/Noggin (WEN)-containing medium devoid of Rspo, forming large cystic structures (**Fig. 5b**), whereas WT organoids did not survive under these conditions.

**Figure 5.**
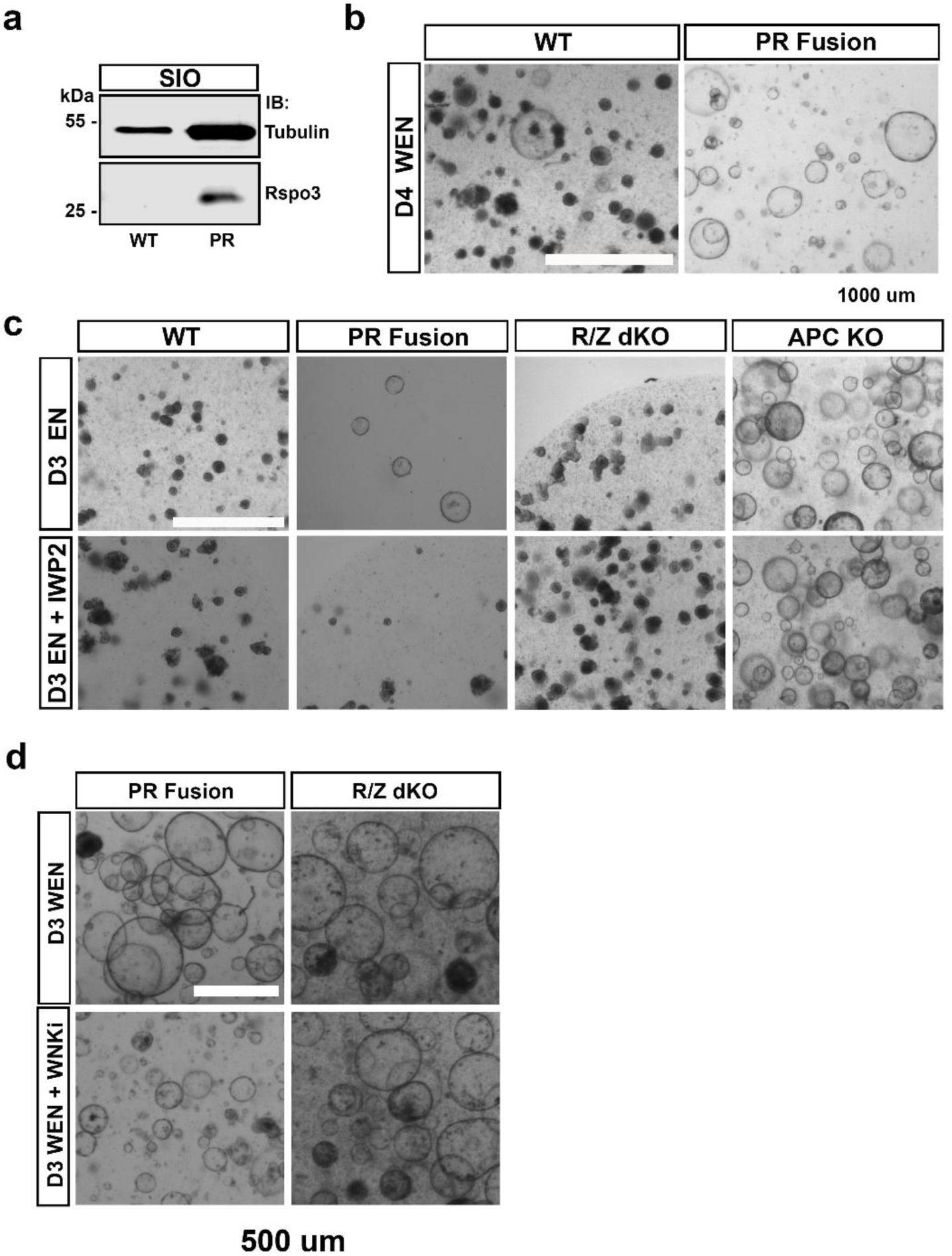
Generation of Ptprk-Rspo3 fusion mouse small intestinal organoids. **a** Western blot showing expression of Rspo3 in Ptprk-Rspo3 mutant organoids (PR). **b** Rspo1 withdrawal assay. At day 4, almost all WT organoids cultured in a medium lacking Rspo1 (EN medium) are dead. However, PR mutant organoids survive Rspo1 deprivation. Scale bar equals 1000 μm. **c** Response of small intestinal organoids with different mutant genetic background to Rspo1 withdrawal, with or without IWP2 Wnt inhibitor. PR fusion organoids survive in the absence of Rspo1, similar to RZ dKO and APC KO mutants. However, PR mutants still depend on endogenous Wnt ligand secretion (like RZ dKO), as treatment with IWP2 kills them. Only APC mutant organoids, which have constitutive Wnt activation, survive IWP2 treatment. WT organoids die in both EN and EN+IWP2 conditions. Scale bar equals 1000 μm. **d** WNKi treatment on PR fusion and RZ dKO organoids. While RZ dKO organoids are insensitive to WNK inhibition, PR mutant organoids stop growing and eventually die in the presence of the WNK inhibitor. Scale bar equals 500 μm.

To further probe their WNT dependency, PR fusion organoids were cultured in EN medium (lacking both Wnt3a and Rspo), with or without the WNT secretion inhibitor IWP2. Similar to Rnf43/Znrf3 double knockout organoids, PR fusion organoids failed to grow when endogenous Wnt secretion was blocked by IWP2, indicating a strong dependency on Wnt ligands (**Fig. 5c**). In contrast, APC KO organoids displayed robust growth in all conditions tested, consistent with ligand-independent, downstream activation of WNT signaling. Finally, treatment with WNK inhibitor greatly reduced the size and number of cystic structures in PR fusion but not R/Z dKO organoids cultured in WEN, suggesting that WNK activity contributes to the growth of a subset of WNT-addicted organoids driven by upstream pathway activation (**Fig. 5d**). These findings identify WNK kinases as a potential therapeutic target in tumors that depend on WNT ligand secretion, such as those harboring RSPO fusions.

## Discussion

Our study reveals a previously unrecognized regulatory axis in canonical WNT signaling involving WNK kinases and the E3 ubiquitin ligase RNF43. While the anti-WNT functions of RNF43 and ZNRF3 have been extensively studied, the upstream modulators of their activity and localization have remained elusive. Through complementary proximity labeling strategies, we identified WNKs as novel interactors of RNF43, linking these kinases to the fine-tuning of WNT receptor turnover at the plasma membrane.

Functionally, we demonstrate that WNK kinase activity negatively regulates the cell surface levels of RNF43, thereby modulating the availability of WNT receptors such as Frizzled and LRP5/6. Pharmacological inhibition of WNK increases RNF43 surface expression, which enhances RNF43-mediated ubiquitination and degradation of WNT receptors, leading to attenuation of canonical WNT signaling. These findings position WNK kinases as critical regulators of WNT receptor homeostasis and expand the mechanistic understanding of how WNT signaling strength is controlled at the receptor level. The interplay between WNKs and RNF43 adds a new layer of complexity to the regulation of WNT signaling. Given that WNKs were traditionally characterized for their roles in ion transport and osmotic regulation, our results broaden their functional repertoire to include modulation of cell signaling pathways relevant for development and cancer. The epistatic relationship between WNKs and RNF43/ZNRF3 suggests that WNKs act upstream of WNT receptor ubiquitination and internalization, potentially by influencing the trafficking machinery or post-translational modifications governing RNF43 stability and localization. Importantly, our data corroborate prior observations linking WNK kinases to β-catenin stabilization and WNT pathway activity (22–24) but provide a more precise molecular context by pinpointing RNF43 as a key mediator. This connection is further supported by the resistance of RNF43/ZNRF3 double knockout cells and organoids to WNK inhibition, highlighting the specificity of WNK action through the RNF43/ZNRF3 axis rather than through downstream canonical effectors.

From a therapeutic standpoint, these findings are particularly relevant. Aberrant WNT signaling is a hallmark of multiple cancers, especially colorectal cancer, where RNF43 mutations are frequently observed (42). Targeting WNK kinases might represent a novel strategy to restore or potentiate RNF43 function, thus re-establishing control over WNT receptor turnover and signaling output. The availability of selective WNK inhibitors enhances the translational potential of this approach, although further studies will be required to elucidate the broader physiological consequences and potential side effects of systemic WNK inhibition. Mechanistically, how exactly WNK kinases modulate RNF43 trafficking remains an open question. Our observations of altered ubiquitination patterns suggest that WNKs may influence the activity of E2 ubiquitin-conjugating enzymes or adaptor proteins involved in endocytosis and lysosomal targeting. The involvement of the WNK downstream kinases OSR1/SPAK in this process also warrants investigation, as they could serve as intermediaries linking WNK signaling to membrane trafficking pathways.

In conclusion, our work uncovers WNK kinases as critical modulators of RNF43 localization and function, adding a novel dimension to the regulation of canonical WNT signaling. This new regulatory axis provides a promising avenue for therapeutic intervention in WNT-addicted tumors and sets the stage for future research into the molecular details and physiological relevance of WNK-mediated control of WNT signaling.

## Material and Methods

### Animals

All animal experiments adhered to the guidelines of the Austrian Animal Care and Use Committee.

### Generation of RNF43 expression vectors

The following plasmids were obtained from Addgene: rat myc-tagged WNK1 (38779); human His-6-,V5-tagged Wnk2 (24569); pCAG-HRP-TM (44441); pcDNA3 APEX2-NES (49386). Plasmids containing human SORTILIN (OHu25285), SORL1 (OHu60668) full length sequences were instead purchased from GenScript. For RNF43-APEX2, a Flag-tagged APEX2 coding sequence was cloned in-frame downstream of RNF43 cytosolic tail, removing the original 2Flag-2HA tag (2F2H) sequence. For HRP-RNF43, the HRP coding sequence was cloned in-frame between the endogenous RNF43 signal peptide and the mature protein sequence. This construct retained the original 2F2H tag previously inserted into RNF43 coding sequence (5). The chimeric constructs were then subcloned into pcDNA4/TO vectors to establish doxycycline-inducible stable cell lines. All constructs reported here were sequence-verified using Sanger sequencing.

### Cell culture, DNA transfection, and growth factor stimulation

HEK 293T cells were maintained in DMEM supplemented with 10% FBS, 1% glutamine, and 1% penicillin-streptomycin, kept in a 37°C and 5% CO_2_ incubator, and passaged every 5 to 7 days. Transfection was performed by using polyethylene imine (PEI; 1 μg/ml), pH 7.4, and plasmid DNA at a 5:1 ratio (16). We transfected 5 μg of total plasmid DNA per 6-cm culture dish (ubiquitination and degradation assays) or 10 μg per 10-cm dish (co-IP experiments). For co-IP experiments, the different construct combinations were transfected at a ratio of 1:1. For Wnt3a treatments, HEK293T cells were plated on 12-well plates, and upon reaching 80% confluency cells were treated with Wnt3a conditioned medium (CM) for 2-3 hours at 37℃. Wnt3a CM was produced from L cells provided by H. Clevers (Hubrecht Institute, Utrecht, Netherlands), following standard protocols. WT L cells (ATCC, CRL-2648) were instead used to produce control CM. Treatments with the WNK inhibitor STOCK2S 26016 were performed at the concentration indicated in the text, overnight at 37℃. DMSO treatments were included as controls.

### Generation of stable cell lines

The pcDNA4/TO HRP-RNF43-2F2H and pcDNA4/TO RNF43-Flag-APEX2 plasmids were transfected into HEK293 T-REx cells using Lipofectamine2000 (Thermo Fisher Sci.). 48 h after transfection zeocin (zeo) was added to the culture medium, at a final concentration of 100 μg/ml. Stable transfected cells were continuously expanded and selected with zeo for at least two weeks, after which monoclonal lines were derived by limiting dilution. Several clones were obtained for each line, which maintained inducible and stable expression of the RNF43 chimeras after many cell passages.

### APEX2- and HRP-mediated proximity labeling

HEK293T-REx stable clonal cells expressing RNF43-APEX2 and HRP-RNF43 were seeded in T225 flasks (Thermo Fisher Sci.) and cultured in filter-sterile growth medium (DMEM supplemented with 10% Fetal Bovine Serum (FBS), 1% penicillin/streptomycin and 1% glutamine), at 37°C and 5% CO_2_. 24 hours before the labeling experiment, approximately at 80% confluency, cells were incubated with doxycycline at a concentration of 1 μg/ml, to induce RNF43 expression. The day after, when cells reached confluency, culture medium was replaced with 25 ml of growth medium supplemented with 500 µM Biotin-Phenol (BP, Iris Biotech). BP was omitted in the no biotinylation (NoBio) control. All samples were incubated with BP for a total of 30 min at 37°C. To start the biotin-labeling reaction, H_2_O_2_ was added directly to the BP-containing medium to a 1 mM final concentration 1 minute before the end of incubation and flasks were gently agitated. At the end of the 1-minute labeling reaction, cells were immediately washed 5 times with quencher solution (ice-cold Dulbecco’s phosphate-buffered saline, DPBS, containing 10 mM sodium azide, 10 mM sodium ascorbate and 5 mM Trolox). To minimize cell detachment during washes, flasks were gently inverted to pour the wash solution on the bottom face, and then inverted again to cover cells with fresh solution. Finally, cells were vigorously washed off the flasks, pelleted by brief centrifugation, and lysed in 1.5 ml of RIPA lysis buffer (50 mM Tris, 150 mM NaCl, 0.1% wt/vol SDS, 0.5% sodium deoxycholate, 1% vol/vol Triton X-100 in nanopure water, pH 7.5, supplemented with 1x Roche cOmplete protease inhibitor cocktail, 10 mM sodium azide, 10 mM sodium ascorbate and 5 mM Trolox). Two T225 flasks were used for each sample, and experiments were conducted in triplicates. For HRP-RNF43 labeling experiment, we followed the same procedure described above with the following modifications: membrane impermeable BP-AEEA (Iris Biotech) was used in place of standard BP, which was added at the 1-min H_2_O_2_-dependent labeling reaction, as described elsewhere (43).

### Mass spectrometry On bead Digestion

Beads were resuspended in 40ul of 100 mM ammonium bicarbonate (ABC), supplemented with 400 ng of lysyl endopeptidase (Lys-C, Fujifilm Wako Pure Chemical Corporation) and incubated for 4 hours on a Thermo-shaker with 1200 rpm at 37°C. The supernatant was transferred to a fresh tube and reduced with 0.5 mM Tris 2-carboxyethyl phosphine hydrochloride (TCEP, Sigma) for 30 minutes at 60°C and alkylated in 4 mM methyl methanethiosulfonate (MMTS, Fluka) for 30 min at room temperature. Subsequently, the sample was digested with 400 ng trypsin (Trypsin Gold, Promega) at 37°C over night. The digest was acidified by addition of trifluoroacetic acid (TFA, Pierce) to 1%. A similar aliquot of each sample was analysed by LC-MS/MS.

### nanoLC-MS/MS Analysis

The nano HPLC system (UltiMate 3000 RSLC nano system, Thermo Fisher Scientific) was coupled to an Exploris 480 mass spectrometer equipped with a FAIMS pro interface and a Nanospray Flex ion source (Thermo Fisher Scientific). Peptides were loaded onto a trap column (PepMap Acclaim C18, 5 mm × 300 μm ID, 5 μm particles, 100 Å pore size, Thermo Fisher Scientific) at a flow rate of 25 μl/min using 0.1% TFA as mobile phase. After loading, the trap column was switched in line with the analytical column (PepMap Acclaim C18, 500 mm × 75 μm ID, 2 μm, 100 Å, Thermo Fisher Scientific). Peptides were eluted using a flow rate of 230 nl/min, starting with the mobile phases 98% A (0.1% formic acid in water) and 2% B (80% acetonitrile, 0.1% formic acid) and linearly increasing to 35% B over the next 120 min. This was followed by a steep gradient to 95%B in 5 min, stayed there for 5 min and ramped down in 2 min to the starting conditions of 98% A and 2% B for equilibration at 30°C. The mass spectrometer was operated in data-dependent mode, performing a full scan (m/z range 350-1200, resolution 60,000, normalized AGC target 100%) at 3 different compensation voltages (CV-45, -60, -75), followed each by MS/MS scans of the most abundant ions for a cycle time of 0.9 (CV -45, -60) or 0.7 (CV -75) seconds per CV. MS/MS spectra were acquired using HCD collision energy of 30%, isolation width of 1.0 m/z, resolution of 30.000, max fill time 100ms, normalized AGC target of 200% and minimum intensity threshold of 2.5E4. Precursor ions selected for fragmentation (include charge state 2-6) were excluded for 45 s. The monoisotopic precursor selection (MIPS) filter and exclude isotopes feature were enabled.

### Proteomics data analysis

Raw MS data was loaded into Proteome Discoverer (PD, version 2.5.0.400, Thermo Scientific). All MS/MS spectra were searched using MSAmanda v2.0.0.16129 (44). Trypsin was specified as a proteolytic enzyme cleaving after lysine and arginine (K and R) without proline restriction, allowing for up to 2 missed cleavages. Mass tolerances were set to ±10 ppm at the precursor and fragment mass level. Peptide and protein identification was performed in two steps. An initial search was performed against the uniprot_reference_human_2022-12-19.fasta (20 520 sequences; 11 403 019 residues) with common contaminants appended. Here, Beta-methylthiolation of cysteine was searched as fixed modification, whereas oxidation of methionine, deamidation of asparagine and glutamine were defined as variable modifications. Results were filtered for a minimum peptide length of 7 amino acids and 1% FDR at the peptide spectrum match (PSM) and the protein level using the Percolator algorithm (45) integrated in Proteome Discoverer. Additionally, an Amanda score of at least 150 was required. A sub-database of proteins identified in this search was generated and used for a second search, where the RAW-files were searched using the same settings as above plus considering additional variable modifications: phosphorylation on serine, threonine and tyrosine, biotinylation on lysine, glutamine to pyro-glutamate conversion at peptide N-terminal glutamine and acetylation on protein N-terminus. The localization of the post-translational modification sites within the peptides was performed with the tool ptmRS, based on the tool phosphoRS (46). Identifications were filtered using the filtering criteria described above, including an additional minimum PSM-count of 2 per protein in at least one sample. The identifications were subjected to label-free quantification using IMP-apQuant (47). Proteins were quantified by summing unique and razor peptides and applying intensity-based absolute quantification (48) with subsequent normalisation based on the MaxLFQ algorithm (49). Identified proteins were filtered to contain at least 3 quantified peptide groups. Statistical significance of differentially expressed proteins was determined using limma (50).

### Volcano Plot Generation

Volcano plots were generated using log2fold changes and −log10 p-values derived from HRP- and APEX2-based biotinylation proteomic experiments. Logarithmic fold changes are plotted on the x-axis, and imputed p-values on the y-axis. Significantly enriched proteins in the HRP-RNF43 and RNF43-APEX2 conditions are presented on the left and right sides of the plot, respectively. Proteins meeting the significance threshold (fold change ≥ 2, p < 0.05) are shown in red. Proteins with fold change ≥ 2 but non-significant p-values are shown in green, while those with p < 0.05 but fold change < 2 are depicted in blue. Data for HRP-RNF43 were generated from triplicate experiments and controls; RNF43-APEX2 analyses used triplicate experiments and duplicate controls. Visualization was performed in RStudio using the EnhancedVolcano package (https://github.com/kevinblighe/EnhancedVolcano).

### Heatmap generation

Heatmaps of proteomic expression for RNF43-HRP and RNF43-APEX2 interactors were generated using row z-scores to normalize protein expression across experimental and control conditions. Row z-scores represent the number of standard deviations a protein’s expression deviates from its mean across all conditions, with red and blue indicating higher and lower expression levels, respectively. HRP-RNF43 heatmaps were derived from triplicate experiments and controls, while RNF43-APEX2 heatmaps were generated from triplicate experiments and duplicate controls. All heatmaps and hierarchical clustering dendrograms were visualized using the heatmap.2 function from the *gplots* package in RStudio publicily available at http://CRAN.R-project.org/package=gplots.

### KEGG Pathway Enrichment Analysis

Lollipop plots were used to visualize pathway enrichment based on the Kyoto Encyclopedia of Genes and Genomes (KEGG) analysis of 577 differentially expressed proteins from the RNF43-HRP experiment and 183 from the RNF43-APEX2 experiment (fold change > 2, p < 0.05). The x-axis represents the number of proteins associated with each pathway, while the y-axis lists the top seven enriched pathways. Color intensity corresponds to protein count (redder shades indicate higher counts), and circle size at the end of each bar reflects statistical significance, calculated as −log10 of the false discovery rate (FDR). Analyses were performed using the clusterProfiler package and visualized with the enrichplot package in RStudio (https://yulab-smu.top/biomedical-knowledge-mining-book/index.html) (51).

### Oncology Subtype Enrichment and UpSet Plot Visualization

Significantly increased HRP-RNF43 proteins (fold change > 2, p < 0.05) were analyzed for disease enrichment using publicly available annotation databases. Proteins were categorized based on their association with cancer-related terms. An UpSet plot was generated to visualize the overlap of enriched proteins across multiple oncology subtypes. Vertical bars represent the number of proteins shared among the cancer categories indicated by connected dots below the x-axis. Isolated dots denote proteins uniquely associated with a single subtype. The y-axis indicates the number of enriched proteins per intersection. This analysis was performed and visualized in RStudio using the UpSetR package available at https://cran.r-project.org/package=UpSetR, which enables scalable visualization of intersecting sets and their distribution across defined categories.

### Gene-Concept Network (cnetplot) Analysis

To explore the biological and oncologic relevance of significantly increased HRP-RNF43 proteins (fold change > 2, *p* < 0.05), a gene-concept network analysis was performed using the **cnetplot** function within the **clusterProfiler** package in RStudio. This visualization displays the associations between enriched cancer-related terms and individual proteins, allowing for the identification of shared nodes across multiple annotations. Proteins are represented as individual dots and are color-scaled by log2 fold change (white to red). Nodes (larger colored circles) represent enriched biological or disease terms, with node size corresponding to the number of associated proteins (cluster size). The proximity of a protein to a node reflects the strength and specificity of the annotation, while proteins linked to multiple nodes indicate shared involvement across several malignant pathways. HRP-RNF43 interactors are indicated in bold to emphasize their functional connectivity within the network.

### Immunofluorescence and confocal imaging

HEK293T cells were grown in 24-well plates containing glass coverslips, precoated with a solution containing 0.01% poly-L-ornithine (Sigma-Aldrich, P4957) overnight at 37°C. Cells were then transfected and/or stimulated as described in text and figure legends, washed twice with PBS (Gibco, 14190-094), fixed in 4% (w/v) paraformaldehyde (PFA) in PBS for 20 min, and then permeabilized with 0.2% (v/v) Triton X-100 in PBS. Coverslips were then washed with PBS, blocked for 1 hour in blocking buffer consisting of 3% (w/v) BSA in PBS at RT, and incubated with primary antibodies in blocking buffer overnight at 4°C. The next day, cells were washed three times with PBS, incubated with secondary antibodies diluted in blocking buffer for 1 hour at RT, and mounted onto glass slides with ProLong Gold antifade reagent containing DAPI (Life Technologies) to stain cell nuclei. For fluorescent staining of protein biotinylation, cells were incubated with 1 μg/ml of NeutrAvidin Texas Red for 1 hour at RT, during the secondary antibody incubation. Imaging was performed using an inverted LSM 880 Airyscan confocal microscope (Carl Zeiss, Jena, Germany) using 405-, 488-, and 561nm lasers for excitation, and a 20× objective (Plan Apochromat ×20/0.8). For scanning, the following parameters were used: unidirectional scanning, averaging number 8, 8-bit depth. Images were acquired with multitracking for each fluorophore and Zeiss ZEN Black Edition software. ZEN Blue Edition software was used for image analysis.

### Small intestinal organoid establishment and maintenance

Small intestinal organoids were established from WT or RZ dKO (5) mice, as reported previously (16). Briefly, mouse small intestinal crypts were isolated by applying Gentle Cell Dissociation Reagent from STEM CELL technologies at room temperature (RT) for 20 min. About 100 to 150 isolated crypts per well were seeded in Matrigel with ENR (Egf, Noggin, and R-spondin1) or WENR (Wnt3a-containing ENR) + nicotinamide (WENR+Nic) culture medium composed of advanced Dulbecco’s modified Eagle’s medium (DMEM)/F12 supplemented with penicillin-streptomycin, 10 mM Hepes (Gibco), GlutaMAX (Gibco), 1× B27 (Life Technologies), 10 mM nicotinamide (MilliporeSigma; used only in WENR), 1.25 mM N-acetyl cysteine (Sigma-Aldrich), mouse Epidermal Growth Factor (mEGF; 50 ng/ml; PeproTech), mNoggin (100 ng/ml; PeproTech), 10% R-spondin1 CM, 50% Wnt3A CM (only in WENR), and 10 mM ROCK inhibitor (Tocris). Established organoids were routinely passaged at 1:3 to 1:5 ratios every week and maintained in a culture medium without ROCK inhibitor. WNKi treatments on intestinal organoids were performed by adding the inhibitor in the culture medium 24 hours after seeding, at the concentrations and for the duration indicated throughout the text. Culture media were refreshed every 2 days, or every day in the case of WNKi. Organoid imaging analysis was performed using a EVOS M7000 microscope (Thermo Scientific).

### Generation of Ptprk-Rspo3 mutant organoids

To induce the chromosome rearrangements leading to Ptprk-Rspo3 fusion, we used CRISPR/Cas9-based gene editing, following the indications and single guide RNA (sgRNA) sequences reported by others (41). WT mouse small intestinal organoids were electroporated with RNP complexes containing Cas9 protein and a pair of sgRNA guides directed against Rspo3 and Ptprk mouse loci (41). Electroporation was performed following a protocol previously established in our lab (52). GFP plasmid was added to the mixture as a control for electroporation efficiency. One week after electroporation, growing organoids were cultured in media deprived of Rspo1, for selection. In such conditions, only organoids with constitutive expression of PTPRK-RSPO3 can survive. Hence, single surviving organoids were manually isolated to generate monoclonal lines and genotyped to confirm the correct chromosomal rearrangement.

### Surface Fzd5 internalization assay

SNAP-tagged Frizzled5 (SNAP-Fzd5) subcellular localization was monitored in WT and WNKi-treated HEK293T cells in the presence and absence of RNF43 coexpression, as previously described (5). SNAP-surface Alexa 549 (NEB) was applied to label surface SNAP-Fzd5 for 15 min at RT in the dark, following the manufacturer’s instructions. Then, Fzd5-labelled cells were either immediately fixed (0 min) or chased for 30 min, before being fixed and processed for confocal imaging. A similar procedure was followed for cells transfected with SNAP-RNF43. Total levels and surface fractions of SNAP-constructs per cell were quantified using ImageJ. The surface levels per cell were determined by subtracting the intracellular fluorescent signal from the total SNAP-construct signal.

### Nano-Glo HiBiT extracellular detection assay

A HiBiT tag was introduced to the N-terminus of V5-Fzd5 and RNF43-2F2H, immediately after the signal peptide, so that the HiBiT tag would be exposed to the extracellular side. HEK293T cells were transiently transfected with HiBiT-Fzd5 or HiBiT-RNF43, and plated 24 hours later onto 96-well flat clear bottom white polystyrene tissue culture treated microplates, at 100,000 cells per well, and cultured in complete DMEM medium as described above. Cells were then incubated with 5 μg/ml WNKi or DMSO (as a control) overnight. The day after, cells were washed once with PBS, and then 100 μl of fresh media were added to each well. The Nano-Glo 2x detection reagent was prepared by diluting the LgBiT protein and its furimazine substrate at 1:100 and 1:50 ratios, respectively, into the detection buffer supplied in the kit. Hence, 100 μl of detection reagent was added to the cells cultured in 100 μl of fresh media, for a 1:1 final dilution. Cells were then incubated for 10 min at RT with gentle shaking, before luciferase detection using a Synergy 2 microplate reader (Biotek). Fzd5 or RNF43 cell surface retention (expressed as %) was calculated using the relative light unit (RLU) readings based on the following equation: percentage retention = (WNKi treatment RLU / No WNKi control RLU) × 100%. Background luminescence was subtracted from all samples using mock-transfected cells as controls. The assay was performed in biological triplicates.

### Cell surface biotin labeling

Cell surface biotinylation was performed as previously described (53). Briefly, HEK293T cells were grown on six-well plates previously coated overnight with a solution (0.1 mg/ml) of poly-L-ornithine (Millipore), an important requirement to prevent cell detachment. Forty-eight hours after transfection with the indicated DNA constructs, cells were transferred on ice and washed twice with ice-cold PBS. Cell surface proteins were then biotinylated by incubation with the cell membrane-impermeable reagent EZ-Link Sulfo-NHS-SS-Biotin (1 mg/ml; Thermo Fisher Scientific) dissolved in PBS for 30 min at 4°C with gentle agitation. Cells were washed three times with ice-cold quenching solution [50 mM glycine in PBS (pH 7.4)] and twice with ice-cold PBS. Cells were kept on ice for the entire length of the labeling protocol. After the final wash, cells were lysed directly on the plates with 300 μl of TNE lysis buffer [tris-NaCl-EDTA, 50 mM tris-HCl (pH 7.4), 150 mM NaCl, 1 mM EDTA, and 1% NP-40] supplemented with protease inhibitors (Roche). Cell lysates were spun at 13,000 rpm at 4°C on a table-top centrifuge to remove cell debris, and then incubated with streptavidin agarose magnetic beads (Thermo Fisher Scientific) for 3 hours at 4°C using a head-over-head rotator to bind biotinylated proteins. An aliquot of the original cell lysate was saved for input control. At the end of the pull-down, beads were extensively washed with TNE buffer at 4°C with rotation, and then heated to 95°C in the presence of 2× Laemmli buffer (Bio-Rad) to elute proteins, followed by SDS–polyacrylamide gel electrophoresis (SDS PAGE)/Western blot analysis. To assess pull-down specificity, a no-biotin control sample was included, and endogenous transferrin receptor was monitored as a negative control.

### Immunoprecipitation

To confirm interactions between RNF43 and the identified targets, we performed IP assays as previously described (16), with some modifications. Briefly, HEK293T cells seeded onto 10-cm plates were transfected with the constructs indicated, using PEI and 10 μg of total DNA. Construct combinations were transfected at a ratio of 1:1. Forty-eight hours after transfection, cells were lysed in 1 ml of IP lysis buffer [10 mM tris-HCl (pH 7.5), 100 mM NaCl, 2 mM EDTA, 1 mM EGTA, 0.5% (v/v) NP-40, and 10% (v/v) glycerol] supplemented with a protease inhibitor cocktail (cOmplete, EDTA-free, Roche). Cell lysates were clarified by centrifugation (13,000 rpm, 4°C), and then precleared with 20 μl of protein A/G agarose magnetic beads (Thermo Fisher Scientific) for 1 hour at 4°C, on a head-over-head rotator. Beads were removed by magnetic separation and cleared lysates were then incubated with 2 μg of primary antibody overnight at 4°C on a head-over-head rotator. The following day, supernatants were incubated with 40 μl of protein A/G magnetic beads for 2 hours at 4°C on rotation, extensively washed in lysis buffer, resuspended in 40 μl of SDS 2x Laemmli buffer (Bio-Rad), and heated for 5 min at 95°C to elute immunocomplexes, followed by analysis through SDS-PAGE and Western blot.

### Western blotting

SDS-PAGE and Western blots were performed using precast gradient gels (Thermo Fisher Scientific), using standard protocols. Blotted nitrocellulose membranes were analyzed using the Li-Cor software Odyssey 3.0. All primary and secondary antibodies were diluted either in tris-buffered saline plus 0.1% Tween 20 (TBST) containing 2.5% (w/v) of Blotting-Grade Blocker (Bio-Rad), or in Li-Cor Intercept (TBS) blocking buffer supplemented with 0.1% Tween 20.

### Ubiquitination assay

HEK293T cells seeded in 6-cm plates and maintained in DMEM with 10% FBS and penicillin-streptomycin were transfected with ubiquitin-Myc-6xHis, and RNF43-2xFlag-2xHA using PEI. Two days after transfection, samples were harvested and lysed with 500 μl of IP lysis buffer (as described above) supplemented with a protease inhibitor cocktail (cOmplete, EDTA-free, Roche) and 10 mM N-ethylmaleimide (NEM, Sigma-Aldrich, E3876). IP and Western blot analysis was then performed as described above.

### *Xenopus* husbandry and embryo microinjection

WT frogs were obtained from the European *Xenopus* Resource Center, UK, and NASCO, USA, and group-housed at the Institute of Molecular Pathology facilities. All animal handling and surgical procedures were carried out adhering to the guidelines of the Austrian Animal Care and Use Committee. In vitro fertilization was performed as previously described (54). Briefly, testes were surgically removed from a male frog anesthetized in 0.03% tricaine methanesulfonate (Sigma-Aldrich, MS222), and a sperm suspension was obtained by crushing each testis in 1 ml of 1× Marc’s modified Ringers [MMR, 0.1 M NaCl, 2.0 mM KCl, 1 mM MgSO4, 2 mM CaCl2, and 5 mM Hepes (pH 7.4)]. Ovulation of female frogs was induced the night before the experiment by injecting 500 IU of human chorionic gonadotropin. On the day of the experiment, frogs were allowed to spontaneously lay eggs in a high-salt solution (1.2× MMR). Laid eggs were collected and fertilized with 200 to 300 μl of sperm suspension. To remove the jelly coat, fertilized eggs were treated with 2% cysteine in 0.1× MMR, pH 7.8, for about 7 min at RT. Dejellied embryos were then cultured in 0.1× MMR’s solution and staged according to Nieuwkoop and Faber (55). For mRNA injection experiments, all RNF43 constructs generated here were subcloned into pCS2+ plasmids. These were then linearized using NotI restriction enzyme and mRNA was transcribed in vitro using mMESSAGE mMACHINE SP6 Transcription Kit (Thermo Fisher Scientific), following manufacturer’s instructions. To induce *Xenopus* double axis, 4 pg of *xWnt8* (Addgene, #16866) mRNA were injected into the equatorial region of a ventral blastomere of four-cell stage embryos. *xWnt8* mRNA was co-injected either with 200 pg of *LacZ* mRNA (for controls) or together with 200 pg of *RNF43* chimeras. After injection, embryos were collected at the early tailbud stage, fixed in 4% PFA in PBS for 2 hours at RT, and washed extensively with PBS to remove residual PFA. Images were acquired on a color camera equipped stereomicroscope (Zeiss).

### Statistics

Statistical analysis was performed using the corresponding functions in Microsoft Excel as well as using R software. Student’s unpaired t tests (two-tailed) were used to compare differences between the two groups. Data from three or more independent groups were analyzed using a two-way analysis of variance (ANOVA). For all statistic tests, *P < 0.05, **P < 0.01, or ***P < 0.001 were considered as thresholds for statistical significance. Unless indicated otherwise in the figure legends, all data presented are mean values ± SD.

## Data availability

All data supporting the findings of this study are available within the Article and its Supplementary Information.

## Additional information

Supplementary information: the online version contains supplementary material available at…

## Acknowledgments

G.C. was supported by the Austrian Science Fund (FWF), Lise Meitner Program M2976 and ERA PerMed (I5900) grants. E.A.S was supported by the Ruth L. Kirschstein Predoctoral Individual National Research Service F31-Award [1F31MH131380-02] and Albert Einstein College of Medicine Medical Scientist Training Program (MSTP) grant [T32-GM149364]. B.-K.K. and his team are supported by the Institute for Basic Science. Proteomics analyses were performed by the Proteomics Facility at IMP/IMBA/GMI using the VBCF instrument pool

## Authors contribution

G.C. designed the project and experiments. G.C., I.J., J.H., S.H.W., K.T., A.C.B. and F.F. performed the experiments. G.C., I.J., A.C.B., N.U., M.M. and B.K. analyzed the data. E.A.S analyzed proteomic enrichment data and performed all associated bioinformatic analyses. E.A.S generated bioinformatic figures related to the HRP-RNF43 and APEX2 datasets, including volcano plots, heatmaps, KEGG pathway enrichment analyses, UpSet plots, and gene-concept network visualizations. G.C. wrote the first draft of the manuscript, which was revised with inputs from I.J., E.A.S., K.T. and B.K.

---

**Supplementary Figure 1.**
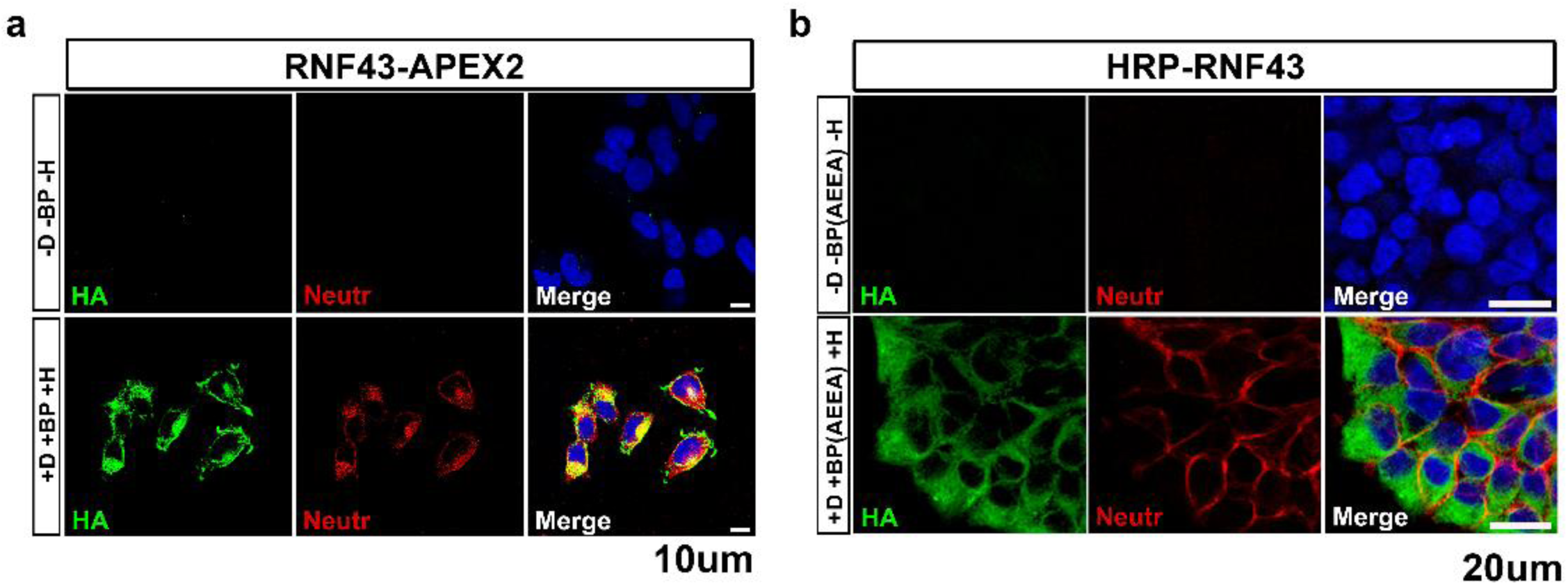
Biotinylation activity in HEK293T cells stably expressing RNF43 chimeric constructs. **a** Neutravidin fluorescent staining reveals strong protein biotinylation in different cytosolic compartments as well as the plasma membrane. Biotinylation is dependent on RNF43 expression (here revealed via HA immunostaining). **b** HRP-RNF43 staining shows strong biotinylation only on the cell surface.

**Supplementary Figure 2.**
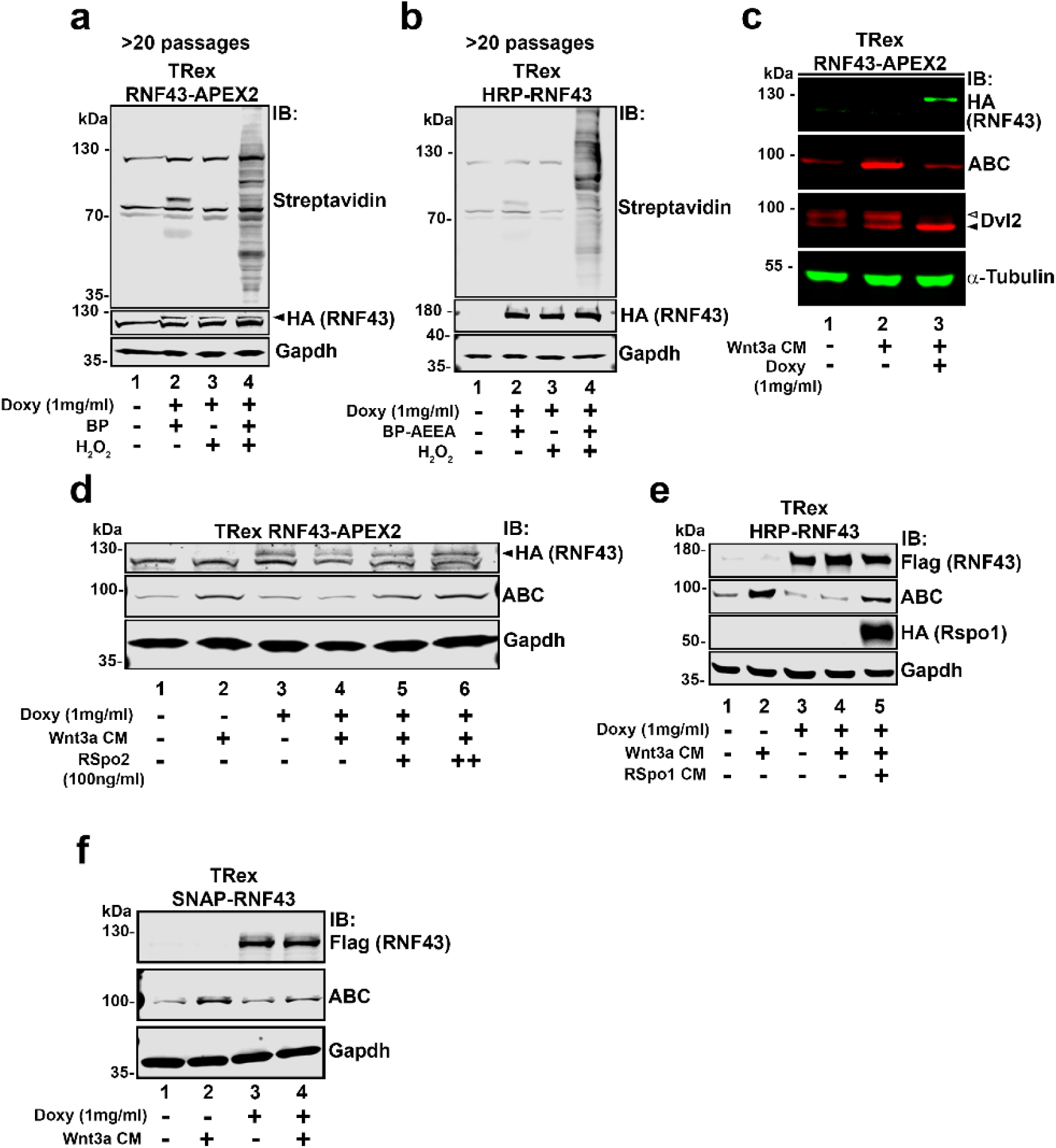
WNT signaling is effectively suppressed by the expression of RNF43-APEX2 and HRP-RNF43. **a, b** Western blot showing doxy-inducible expression of RNF43-APEX2 and HRP-RNF43 in stably transfected cells, even after many passages. **c** Western blot showing that, in doxycycline treated cells, RNF43-APEX2 strongly reduces active β-catenin (ABC) and phospho-Dvl2 (white arrowhead; black arrowhead indicates non-phosphorylated Dvl2) levels, as expected. **d, e** Western blot experiments confirm that both RNF43-APEX2 and HRP-RNF43 maintain not only WNT inhibitory activity, as assessed by reduction of ABC, but also responsiveness to R-spondin ligands, which relieve RNF43-mediated WNT suppression. **f** Western blot showing that another recombinant RNF43 construct, SNAP-RNF43, maintains WNT inhibitory activity, as determined by reduction of ABC protein levels.

**Supplementary Figure 3.**
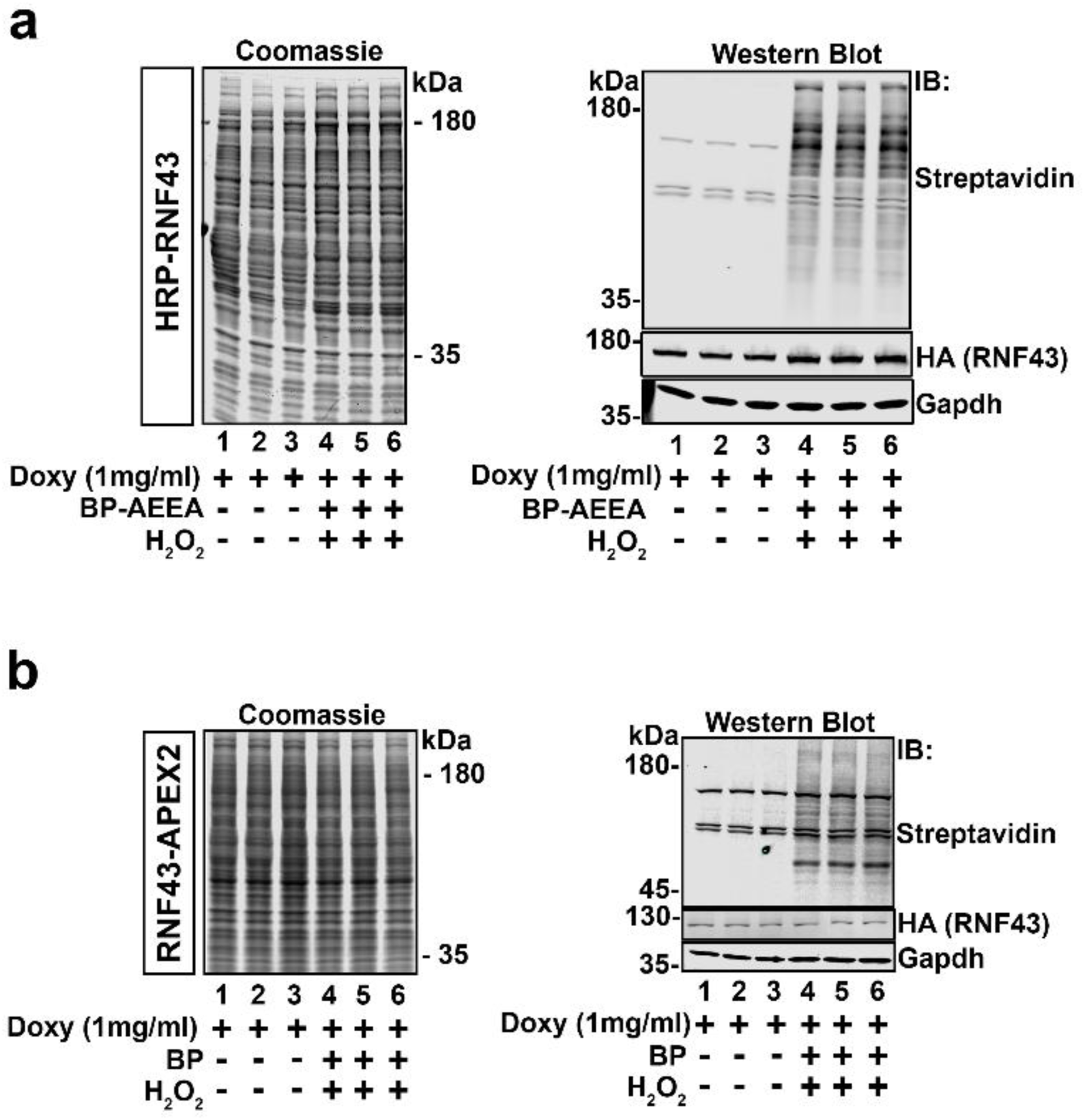
Quality control blots for mass spectrometry analysis. **a, b** Blue coomassie gel staining showing equal amount of protein loading for each sample (3 controls, lanes 1, 2 and 3; 3 treated, lanes 4, 5 and 6). Western blots confirm that biotinylation of RNF43 interactors only occurs in the treated samples (lanes 4, 5 and 6).

**Supplementary Figure 4.**
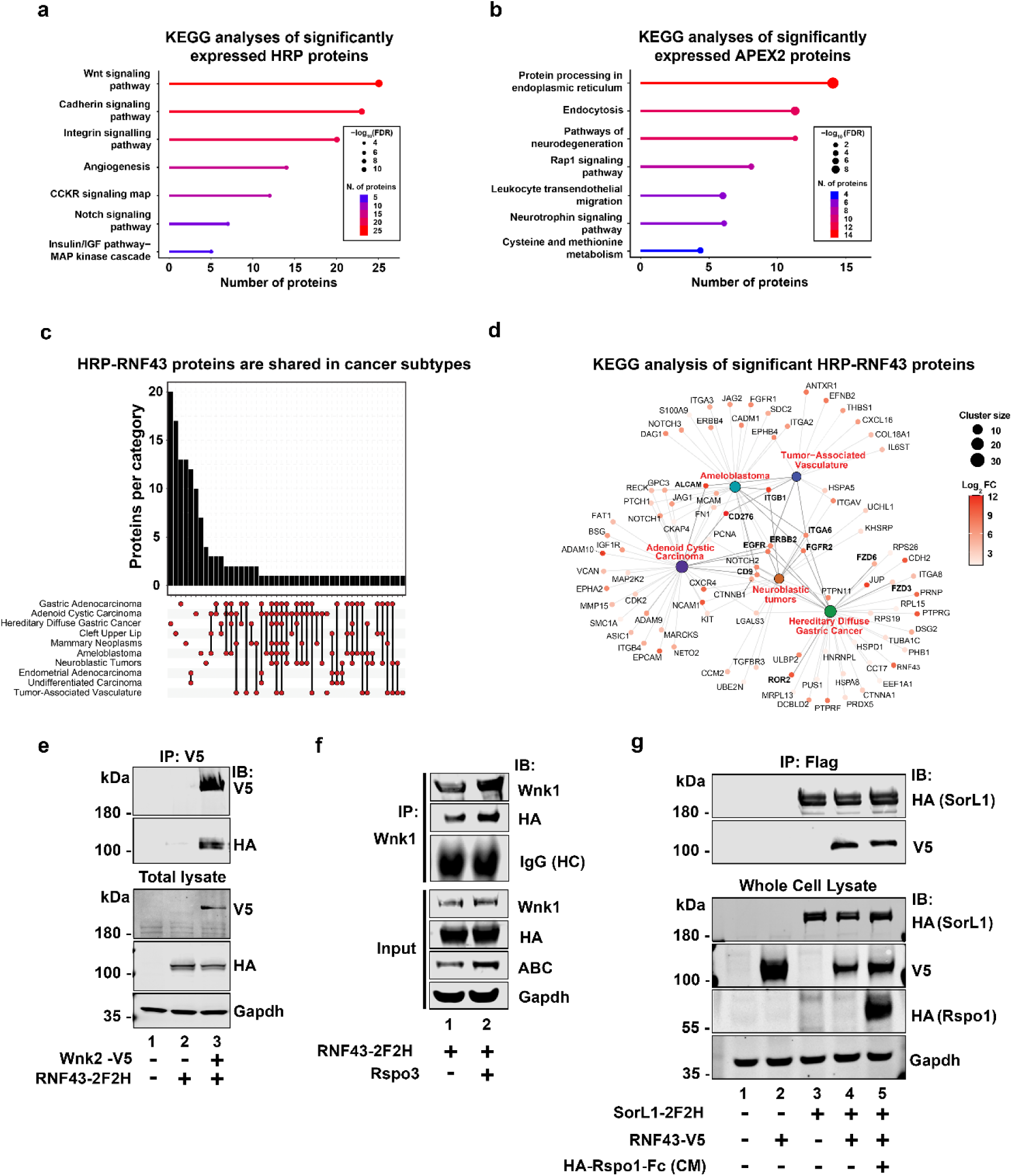
Bioinformatic analysis of the RNF43 interactome. **a** Lollipop plot of KEGG pathway enrichment among the 577 differentially expressed HRP-RNF43 proteins (fold change ≥ 2, *p* ≤ 0.05). The x-axis denotes the number of associated proteins per pathway; circle size reflects statistical significance (−log_10_FDR). The top seven pathways are shown, with Wnt signaling most enriched, followed by Cadherin and Integrin signaling. **b** KEGG pathway analysis of 183 RNF43-APEX2 differentially expressed proteins (fold change ≥ 2, *p* ≤ 0.05), highlighting the top seven pathways. Protein processing in the endoplasmic reticulum is the most significantly enriched. **c** UpSet plot of HRP-RNF43 proteins across oncology subtypes using only significant proteins (fold change ≥ 2, *p* ≤ 0.05). Red dots without connecting lines indicate disease-specific proteins, while connected dots denote overlap across multiple cancer types. Vertical line length corresponds to the number of shared subtypes. The y-axis indicates the number of proteins per cancer category. **d** Gene-concept network (cnetplot) illustrating connections between significant HRP-RNF43 proteins and malignancy-related pathways. Proteins are represented as dots, colored by fold change (white to red), with bolded labels indicating HRP-RNF43 interactors. Node proximity reflects strength of literature-based association, and node size indicates the number of associated proteins (cluster size). **e, f** IP experiments followed by Western blot analysis showing that RNF43 interacts with overexpressed WNK2 (**e**) or endogenous WNK1 (**f**). Note that Rspo1 treatment does not increase RNF43/WNK interaction. **g** IP experiment followed by Western blot analysis confirming RNF43 interaction with Sortilin-like1 (SorL1) protein. Treatment with Rspo1 does not have any effect on this interaction (lane 4).

**Supplementary Figure 5.**
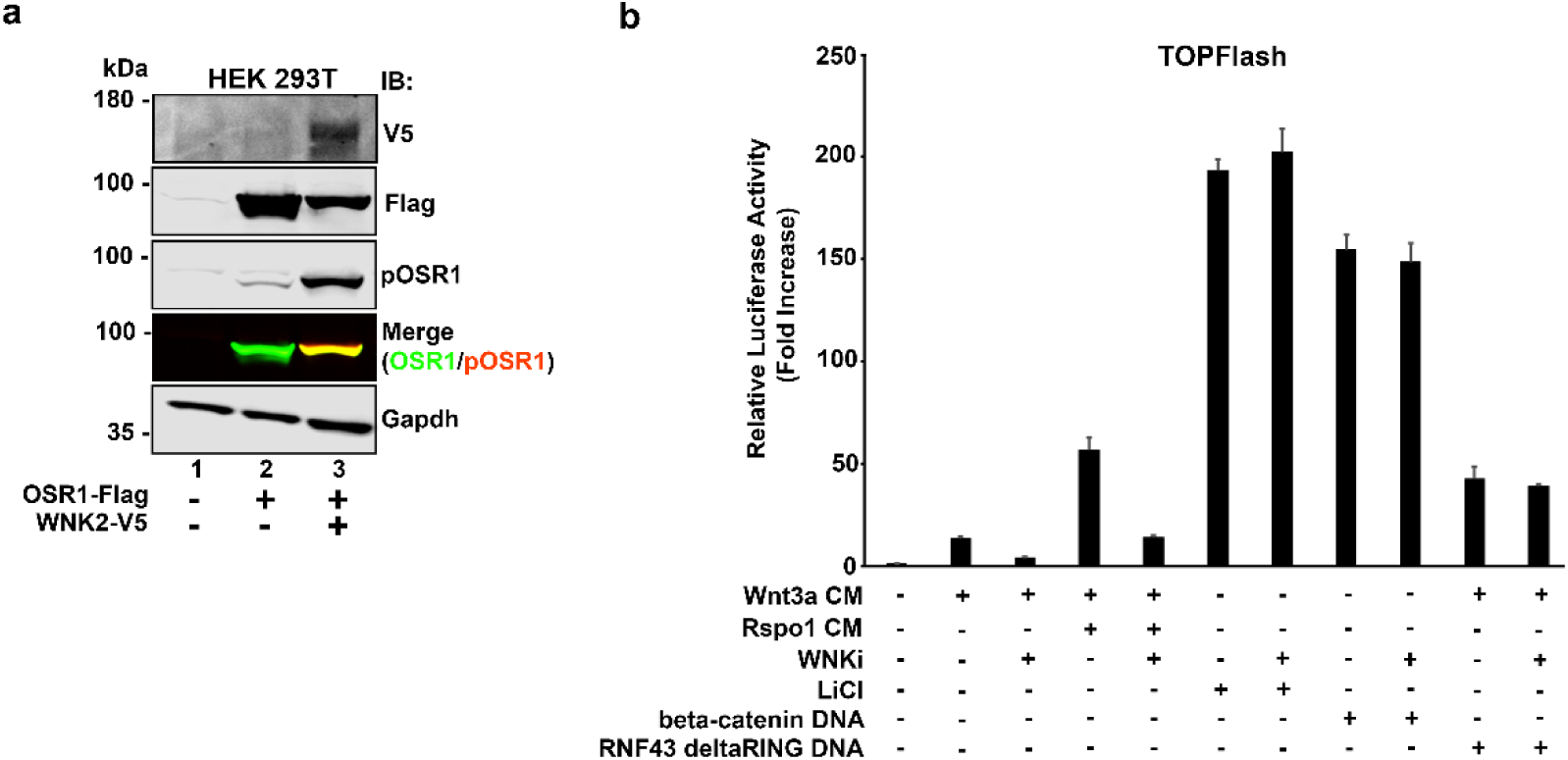
WNK inhibition negatively affects WNT signaling. **a** Western blot showing that overexpression of WNK is sufficient to phosphorylate and activate its substrate OSR1, thus stimulating the WNK cascade. **b** WNT reporter TOPFlash analysis showing that WNK inhibition (by the chemical compound STOCK2S-26016, or WNKi) strongly reduces WNT stimulation by Wnt3a and Rspo1 ligands. However, note that WNKi does not have any effect on WNT induction by overexpressed β-catenin, dominant negative RNF43 or upon treatment with the GSK3 inhibitor, LiCl. This suggests that WNK inhibitor impairs WNT signaling at the ligand/receptor levels, upstream of β-catenin and the β-catenin destruction complex.

**Supplementary Figure 6.**
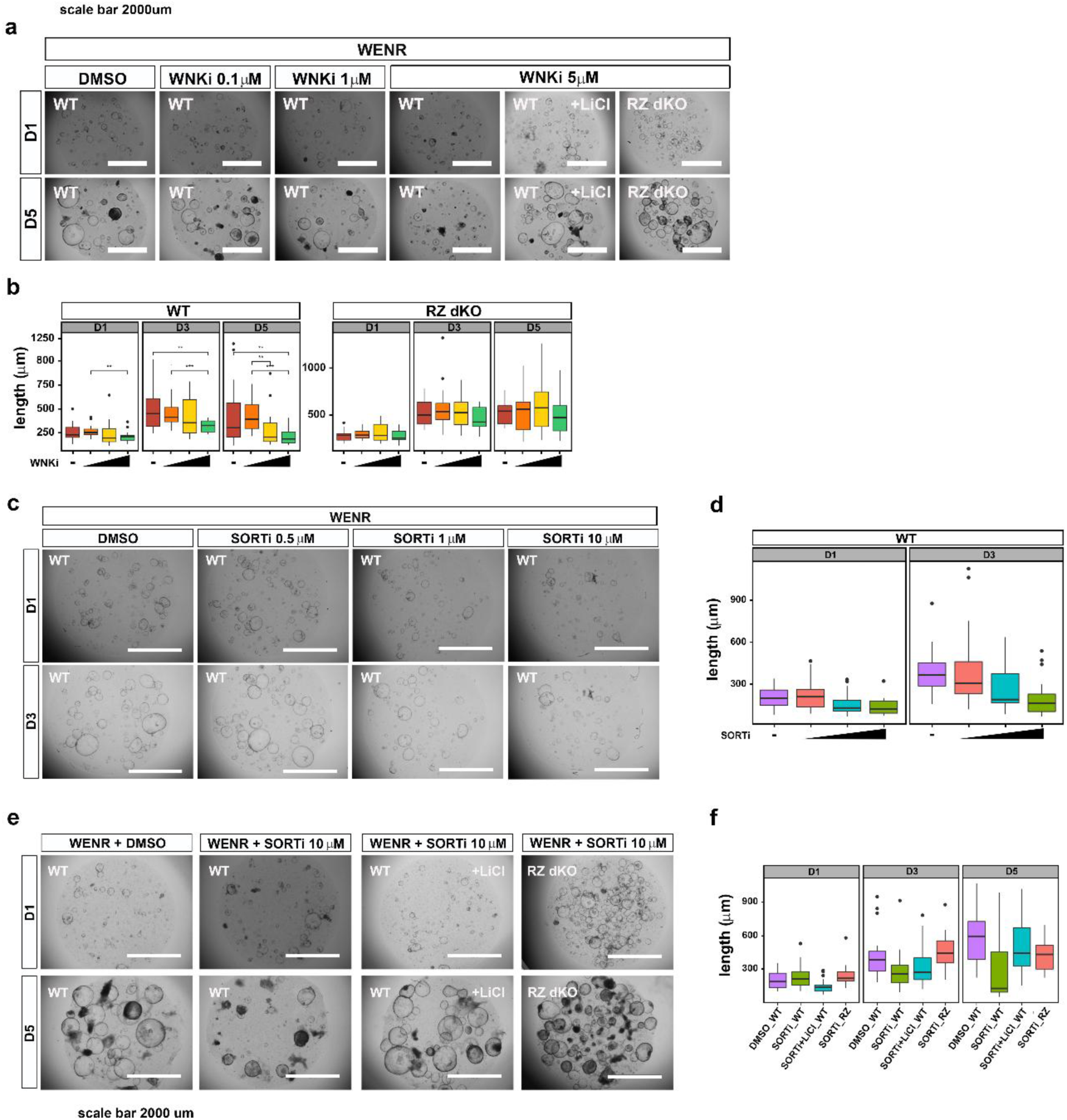
Dose dependent effects of WNK and Sortilin inhibitors on mouse small intestinal organoids. **a** WT organoids growth is progressively reduced by higher concentrations of WNKi. Note that even the highest concentration of WNKi used in this study, 5 μM, has no effects on the growth of RZ dKO organoids. **b** Quantification of the experiment shown in **a**. **c** Similarly to WNKi, Sortilin inhibition also reduces organoid growth in a dose-dependent fashion, albeit the effects are milder compared to WNK inhibition. **d** Quantification of the experiment shown in **c**. **e** While Sortilin inhibition inhibits WT organoids growth, it does not have any effect on LiCl treated or RZ dKO organoids. **f** Quantification of the experiment shown in **e**.

